# Higher throughput assays for understanding the pathogenicity of variants of unknown significance (VUS) in the RPE65 gene

**DOI:** 10.1101/2025.01.31.635952

**Authors:** Leila Azizzadeh Pormehr, Kannan Vrindavan Manian, Ha Eun Cho, Jason Comander

## Abstract

**Purpose:** *RPE65* is a key enzyme in the visual cycle that regenerates 11-cis retinal. Mutations in *RPE65* cause a retinal dystrophy that is treatable with an FDA-approved gene therapy. Variants of unknown significance (VUS) on genetic testing can prevent patients from obtaining a firm genetic diagnosis and accessing gene therapy. Since most *RPE65* mutations have a low protein expression level, this study developed and validated multiple methods for assessing the expression level of *RPE65* variants. This functional evidence is expected to aid in reclassifying *RPE65* VUS as pathogenic, which in turn can broaden the application of gene therapy for *RPE65* patients.

**Methods:** 30 different variants of *RPE65* (12 pathogenic, 13 VUS, 5 benign) were cloned into lentiviral expression vectors. Protein expression levels were measured after transient transfection or in stable cell lines, using Western blots and immunostaining with flow cytometry. Then, a pooled, high throughput, fluorescence-activated cell sorting (FACS) assay with an NGS-based sequencing readout was used to assay pools of *RPE65* variants.

**Results:** There was a high correlation between protein levels measured by Western blot, flow cytometry, and the pooled FACS assay. Using these assays, we confirm and extend *RPE65* variant data, including that Pro111Ser has a low, pathogenic expression level. There was a high correlation between RPE65 expression and previously reported enzyme activity levels; further development of a high throughput enzymatic activity assay would complement this expression data.

**Conclusion:** This scalable approach can be used to solve patient pedigrees with VUS in *RPE65*, facilitating treatment and providing *RPE65* structure-function information.

## Introduction

One of the primary goals of human genetics is to understand how genetic variation affects the function of genes and contributes to disease development.^1^ In fact, of more than 4 million missense identified variants, only about 2% have been definitively classified as pathogenic or benign. Most missense variants are classified as VUS.^2^ The current study is motivated by the observation that genetic testing for inherited retinal diseases (IRDs) gives ambiguous or negative results about one third of the time, often due to VUS.^3–7^ Proper pathogenicity classification of new or rare variants is important for a conclusive molecular diagnosis and the medical management of patients.^8^ This problem can have clinical consequences, preventing familial risk assessment, family planning, and access to an approved gene therapy or other investigational gene-specific therapies. In IRDs, this problem of VUS is particularly impactful for the *RPE65* gene, which was the first gene to have a corresponding FDA-approved gene therapy. Indeed, genetic testing results containing a *RPE65* VUS can prevent patients from obtaining treatment with an FDA-approved gene therapy, Voretigene neparvovec (Luxturna, Spark Therapeutics, Philadelphia, PA). ^9,10^ Mutations in the *RPE65* gene account for 0.6–6% of RP and 3–16% of LCA/EORD cases,^11^ and patients are eligible for treatment only if they have documented biallelic pathogenic or likely pathogenic mutations. ^12^

While there are a number of computational tools that can be used to predict pathogenicity^2,13,14^, they are not highly accurate.^15,16^ Even with advances in computational algorithms^17–19^, the level of accuracy may not be high enough when making medical decisions, including the decision to expose a patient to a gene therapy surgery specific to a certain genetic cause of disease. Additionally, ACMG guidelines do not allow a definitive diagnosis based on computational predictions alone.^20^ As a result, the use of validated, laboratory-based functional assays is considered strong evidence towards the reclassification of VUS into pathogenic variants.^2,6,7,13–20^

While the traditional method of investigating variant pathogenicity is to test one variant at a time, the benefits of producing this information at scale have resulted in the field of “functional genomics” in which parallelized, higher-throughput assays are used to assay pools of variants. Widespread use of functional genomics could improve the accuracy of variant interpretation in genes with both known and unknown associations with disease, generate information and reagents needed to test therapeutic agents, and inform the development of analytical tools for predicting variant pathogenicity. Therefore, the purpose of this study is to develop and validate higher-throughput expression assays for *RPE65* to provide expression information on a panel of *RPE65* variants at a higher accuracy than can be achieved by bioinformatic estimates alone.

Mechanistically, *RPE65* plays the central role in the retinoid cycle^21–23^ and encodes retinoid isomerohydrolase. The retinoid cycle enzymatic pathway allows continuous vision by regeneration of 11-cis from all-trans-retinal, which becomes part of the main chromophore of phototransduction in photoreceptor cells.^24–26^ More than 230 missense mutations lacking a clear pathogenicity classification or classified as VUS have been reported in public databases for *RPE65* (ClinVar). Many of these mutations introduce missense and nonsense mutations which affect the protein expression level or enzymatic activity, including via changing protein localization or stability.^27–29^ For example, missense mutation of G40S caused reduced catalytic activity to 2% of wildtype levels, and reduced protein levels to less than 40% of wild type levels.^23,30,31^

A traditional method for quantifying protein levels in cells is the Western blot, which is semi-quantitative when calibrated, but is low-throughput and has a limited dynamic range. More scalable quantification techniques can include use of fluorescent protein fusions (VAMP-seq)^32^, flow cytometry^33^, split luciferase systems^34^, and barcoded transcriptional reporters.^35^ Flow cytometry is particularly suited to higher-throughput assays implemented as pooled assays, as individual cells from a library can be rapidly separated for analysis. Fluorescence from immunostaining or from engineered fluorescent proteins can be used to sort cells based on the expression of cell surface and/or intracellular proteins. By combining FACS and next-generation sequencing (NGS), it is possible to obtain a more comprehensive understanding of thousands of variants.^36^ While many functional genomics studies use tags to facilitate protein detection ^37,38^, this study uses only the native to human *RPE65* protein sequence to reduce the uncertainty that genetically encoded tags could disrupt the fidelity of the assays in a manner that is difficult to detect.

We hypothesized that cross-validating different analytical techniques with different antibodies and across a range of *RPE65* variants would evaluate the specificity and reproducibility of the reagents and allow for the development of a higher throughput RPE65 protein expression assay based on flow cytometry.

## Methods

### Selection of *RPE65* variants for study

30 different *RPE65* variants were selected spanning a variety of pathogenicity levels: 12 pathogenic, 13 VUS, 5 benign. (See Supplemental Table 1, with HGVS nomenclature.) Wildtype RPE65 was also included. Among the variants that underwent analysis by multiple assays (Table 1), the negative control mutants that are known to have low expression levels included: G40S, R515W, G104V, P25L, P363T, G244V, and R124*. Y368H has conflicting expression level data and is known to have low activity. H241L and Y239D were selected as known mutations in the active site region. C112Y was reported as homozygous in IRD pedigrees ^39,40^, which this study will consider as likely pathogenic. The positive control variants were A434V and N321K, which are known to have normal ^30^ or slightly decreased ^23^ activity and have a high maximum allele frequency in humans: A434V: 7.7% in African/African Americans and N321K: 3.5% in South Asians (gnomAD database v4.1.0). K294T was a likely benign variant, with somewhat conflicting activity levels reported^23,30^. The variants N248S, V189I, T94A, T390I were selected from a panel of VUS that were detected in IRD patients, provided by Spark Therapeutics.

**Table 1.**
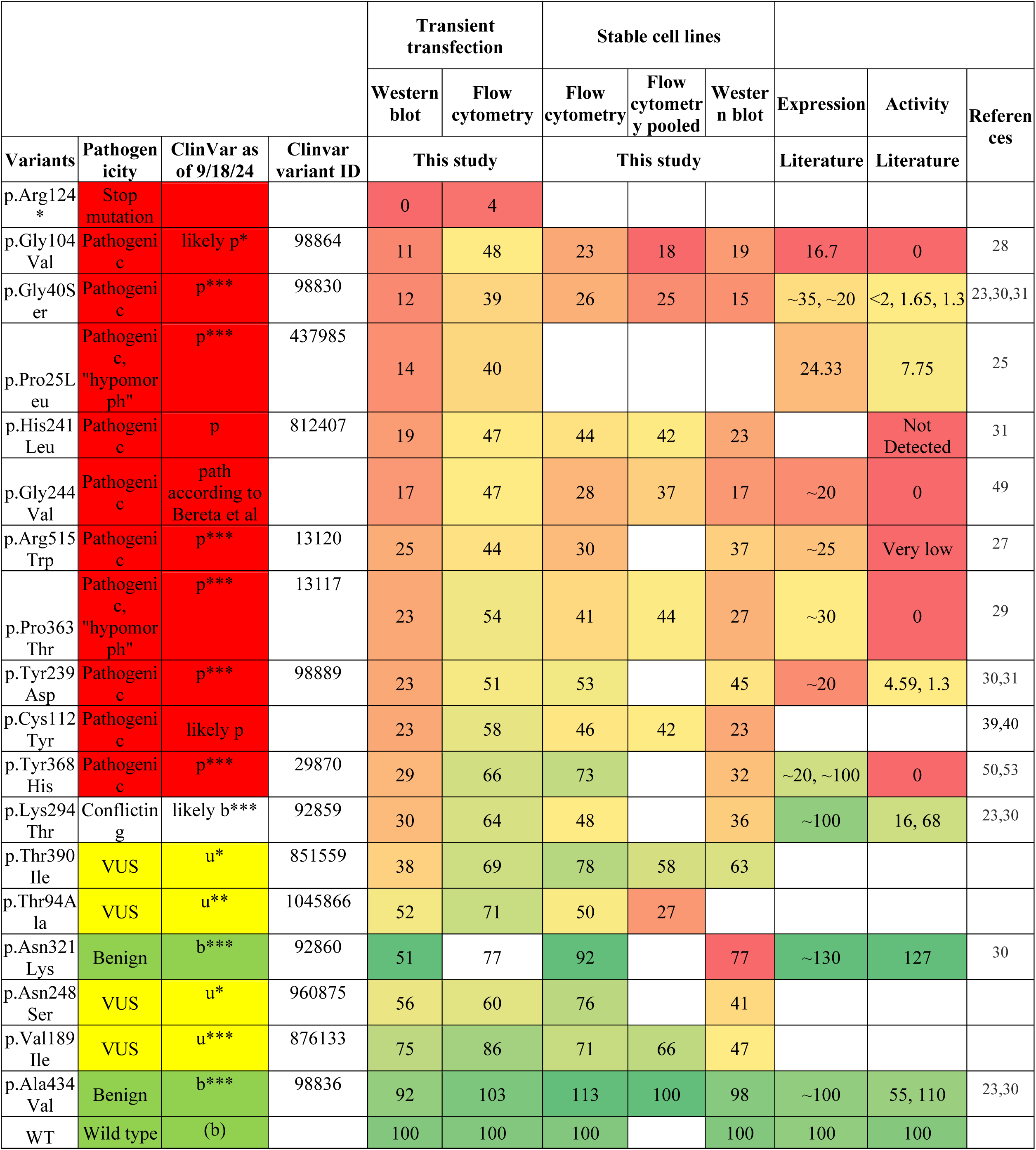
Comparing expression levels of different known RPE65 variants between the results in this study and in previous studies. Results are expressed as a percent of wildtype. The color code for values is green=100 and red =0. The color code for pathogenicity is green = benign (b), yellow = VUS (u), red=pathogenic (p). Asterisks denote the number of stars in the ClinVar evidence rating.

A second batch of variants were selected for testing by flow cytometry analysis only (Table 2). A434=, T385=, and E352= are synonymous variants hypothesized to behave as wildtype. T86N, S533T, T105N, P111S, and N135K was selected because, at the time of initiation of this project, the variants were identified as VUS by the RPE65 Variant Curation Expert Panel (Lori Sullivan; personal communication). G193S was selected as a VUS from the ClinVar database. L450V is a potential hypomorphic allele from our institution’s patient cohort. D477G is a known dominant pathogenic mutation which produces a distinct phenotype compared to recessive *RPE65* mutations ^23^. H527R is a known mutation in the active site region.

**Table 2.**
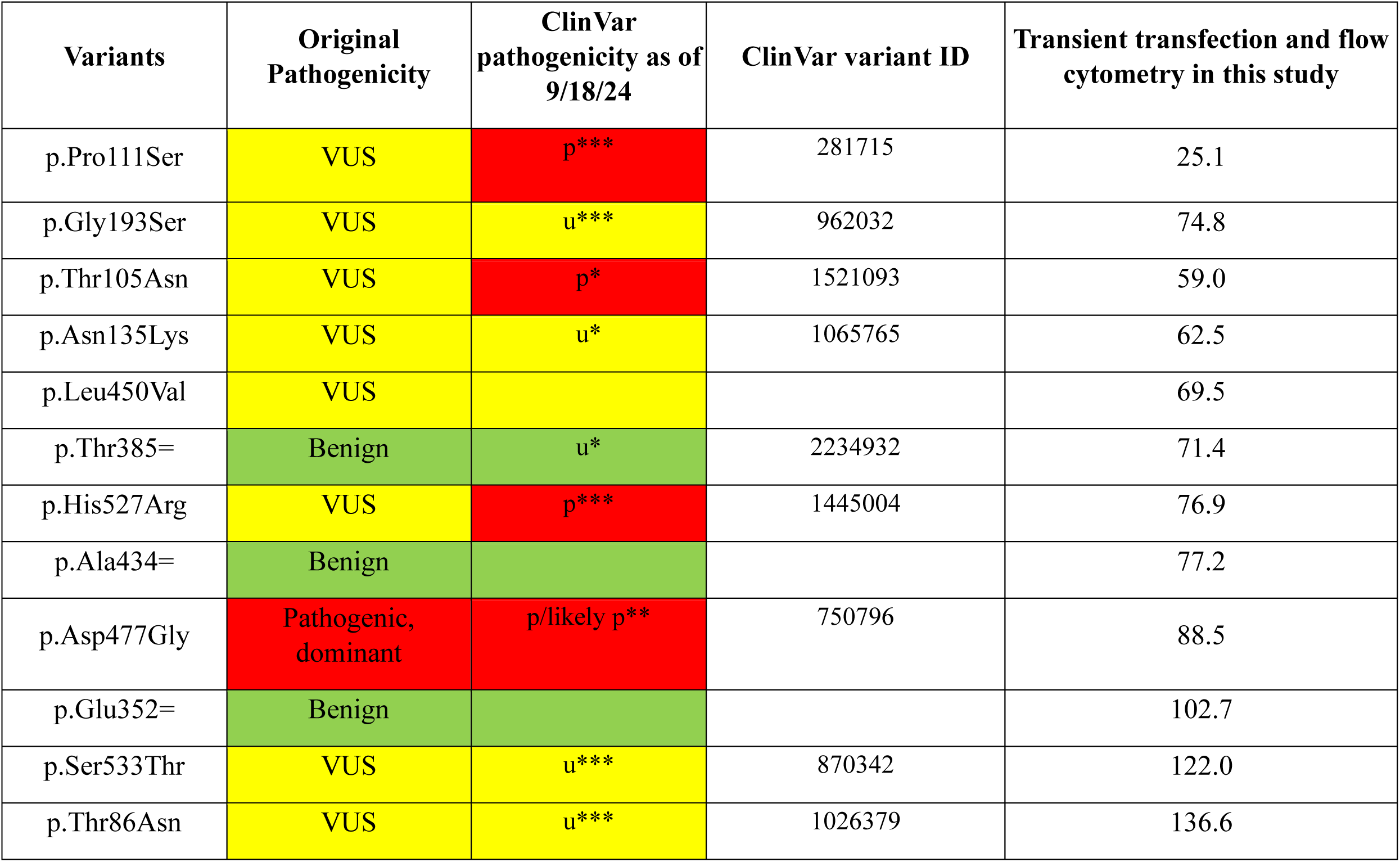
Expression levels of additional RPE65 variants that were assay using the transient transfection method. The color code for pathogenicity is green = benign (b), yellow = VUS (u), red=pathogenic (p). Asterisks denote the number of stars in the ClinVar evidence rating.

### Plasmid construction

The human *RPE65* cDNA (NM_000329.3, without untranslated regions) amplified and was cloned into the NheI and BamH1 site of pMT_025 lentiviral expression vector (Addgene,#158579).^41^ Mutagenesis was used to create the DNA changes as listed in **Supplementary Table 1**. NGS or Sanger sequencing was performed to verify the correct sequence in all plasmids.

### Transient transfection of variants

30 different variants of *RPE65* were used for transfection in HEK293T cells (ATCC, Cat no. CRL-3216). For transfection, HEK293T cells were seeded in 6-well(C6) or 12-well(C12) plates at a density of 4.3×10^4^ cells/cm^2^. The cells were maintained in Dulbeccòs Modified Eagle Medium (DMEM) supplemented with 10% fetal bovine serum (Thermo Fisher, Cat no. SH3007103). At 70-80% confluency, 2.5 μg plasmids were transfected per C6 well using Lipofectamine 3000 transfection reagent (Thermo Fisher, Cat no. L3000008) according to the manufacturer’s instructions. Cells were assayed 48 hours post-transfection.

### Generation of stable cell lines for *RPE65* variants

Lentiviruses were prepared by transfecting HEK293FT cells with psPAX2 (Addgene no. 12260), pMD2.G (Addgene no. 12259) and pMT_025 RPE65 variant using Lipofectamine LTX according to the manufacturer’s instructions. The viral supernatant was harvested, concentrated and transduced into HEK293T cells at MOI of <0.3. The transduced cells were selected using puromycin to generate stable lines. For additional details, see Supplementary Methods.

### Western blotting

Whole-cell lysates were prepared using RIPA buffer from the transient and stable cell lines. After quantification, the lysates were loaded on SDS-PAGE, followed by dry transfer using iBlot™ 2 system (Invitrogen, Carlsbad, CA). Blots were incubated with primary antibodies of *RPE65* and ß-actin overnight at 4°C followed by respective secondary antibodies. The blots were detected using the Odyssey CLx infrared imaging system (LICOR, Lincoln, Nebraska). LI-COR Image Studio was used to quantify the band intensity, and the data was normalized to the expression level seen in the wild-type (WT) *RPE65* sample. For additional details, see Supplementary Methods.

#### Quantification of RPE65 expression levels by flow cytometry

Initial experiments used the PETLET *RPE65* antibody (not shown; from the Redmond laboratory), but the following experiments were conducted with commercially available antibodies that are more broadly available. Cells were collected for flow cytometry analysis 48 hours post transfection. For stable cells lines, cells were collected at 80% confluency. After washing with PBS, cells were collected by trypsinization, centrifuged at 1200 g for 4 minutes, and washed with PBS. The cells were resuspended and fixed in 4% paraformaldehyde (PFA, Electron Microscopy Sciences, Cat no. 15714) in PBS for 20 min and washed with PBS and kept at 4^0^C until staining. For staining of *RPE65*, cells were permeabilized with 0.02% saponin (Sigma, Cat no. 47036) in PBS for 12 minutes, followed by blocking in 3% BSA in PBS. After blocking, cells were incubated with primary antibodies described above at a final concentration of 1:1500 in blocking buffer for 1 hour. After washing with PBS, Alexa fluor-488 secondary antibodies goat anti-mouse IgG1 (Invitrogen, Cat no. A21121) and goat anti-rabbit IgG (Invitrogen, Cat no. A27034) were used at a final concentration of 1:500 in blocking buffer for 1 hour. After a PBS wash, cells were resuspended in appropriate volume of PBS and analyzed on a Miltenyi BioTec MACSQuant flow cytometer. Flow cytometry results were analyzed with FlowJo Software v10. The resulting data were averaged across multiple independent transfections (N= 3-5).

#### Pooled quantification of *RPE65* expression levels by FACS and NGS

Stable cell lines expressing RPE65 variants (N=16 variants) were pooled together, fixed using 4% PFA, and stained for FACS as described previously for flow cytometry. The stained cells were sorted on Sony SH800 or MA900 FACS machines by their *RPE65* expression level, where the “high” and “low” gates contained the top and bottom 18% of fluorescent intensity, respectively. The cells were lysed and the genomic DNA from *RPE65*^high^, *RPE65*^low^, and unsorted cells was un-crosslinked (manuscript in preparation) and extracted. The open reading frame of the lentiviral integrands was amplified and quantified using next-generation sequencing and compared to flow cytometry expression results. Briefly, the open reading frame containing the *RPE65* variants was amplified using primers XY304 (ATTCTCCTTGGAATTTGCCCTTT) and XY305 (CATAGCGTAAAAGGAGCAACA). NGS libraries (170-280 bp) were created using fragmentation and sequenced on the Illumina MiSeq. Fastq files were aligned to a reference sequence containing the pMT025-RPE65 plasmid. Variants were quantified using bam-readcount^42^ and normalized to read depth. The relative expression level for each variant in the pool was calculated as the variant count in the high gate divided by the count in the low gate.^33^

## Results

### Assay development

#### Measuring the linearity of western blot detection of *RPE65* and Beta-actin

To calibrate the detection range and linearity of *RPE65* and ß-actin via Western blotting, we prepared HEK293T cell lysates transiently transfected with *RPE65*-wildtype (WT) and loaded 0.1 −20 μg of whole cell lysates for immunoblotting (N=2). Densitometry showed that the linear quantification range was narrow and required using relatively small amounts of protein (0.25-2 μg total lysate) and a small amount of antibody (1:5000 dilution of anti-RPE65 EPR), as seen in **Figure 1 A, B**. There was no *RPE65* specific band (65 kDa) observed in untransfected cell lysate (not shown and also see **Figure 2** below). To further delineate the proportional linear range, even smaller amounts of diluted protein lysate (0.05-4 μg) were used, which indicated that the proportional linear range for both *RPE65* and ß-actin (Santa Cruz) were 0.1-2 μg (**Figure 1 C, D**). For stable cell lines that contained only a single copy of the *RPE65* transgene, there was an approximately 4-fold lower expression in the stable lines compared to transient transfection, and protein lysate ranging from 1-20 μg was tested. Four μg was the higher end of linear range (not shown). For further analysis of the different variants by Western blot, 1 μg protein lysate for transient transfections and 4 μg protein lysate for stable cell lines was used per lane. Total protein staining (Revert 700) was not accurately detectable by Li-cor Odyssey densitometry using the small protein amounts required for antibody detection within the linear range (not shown), therefore ß-actin was used as a loading control for normalization. For the single-copy stable cell lines only, the use of 4 μg protein lysate was slightly beyond the linear range of ß-actin detection but was required for detection of *RPE65*. The use of 1 μg protein lysate from the transient transfections was within the linear detection range for both *RPE65* EPR and ß-actin. Similar results were obtained with the “3D9” *RPE65* antibody, but for Western blotting the linear range and detection limit were slightly lower (not shown). Altogether, this data indicated that the best combination of antibodies and conditions for the most accurate quantification of *RPE65* protein levels by Western blot was “EPR” (1:5000) and Santa Cruz sc-47778 anti-ß-actin (1:5000). (See Methods.) Therefore, these antibodies and concentrations were used for comparing *RPE65* variant expression levels using Western blotting below.

**Figure 1.**
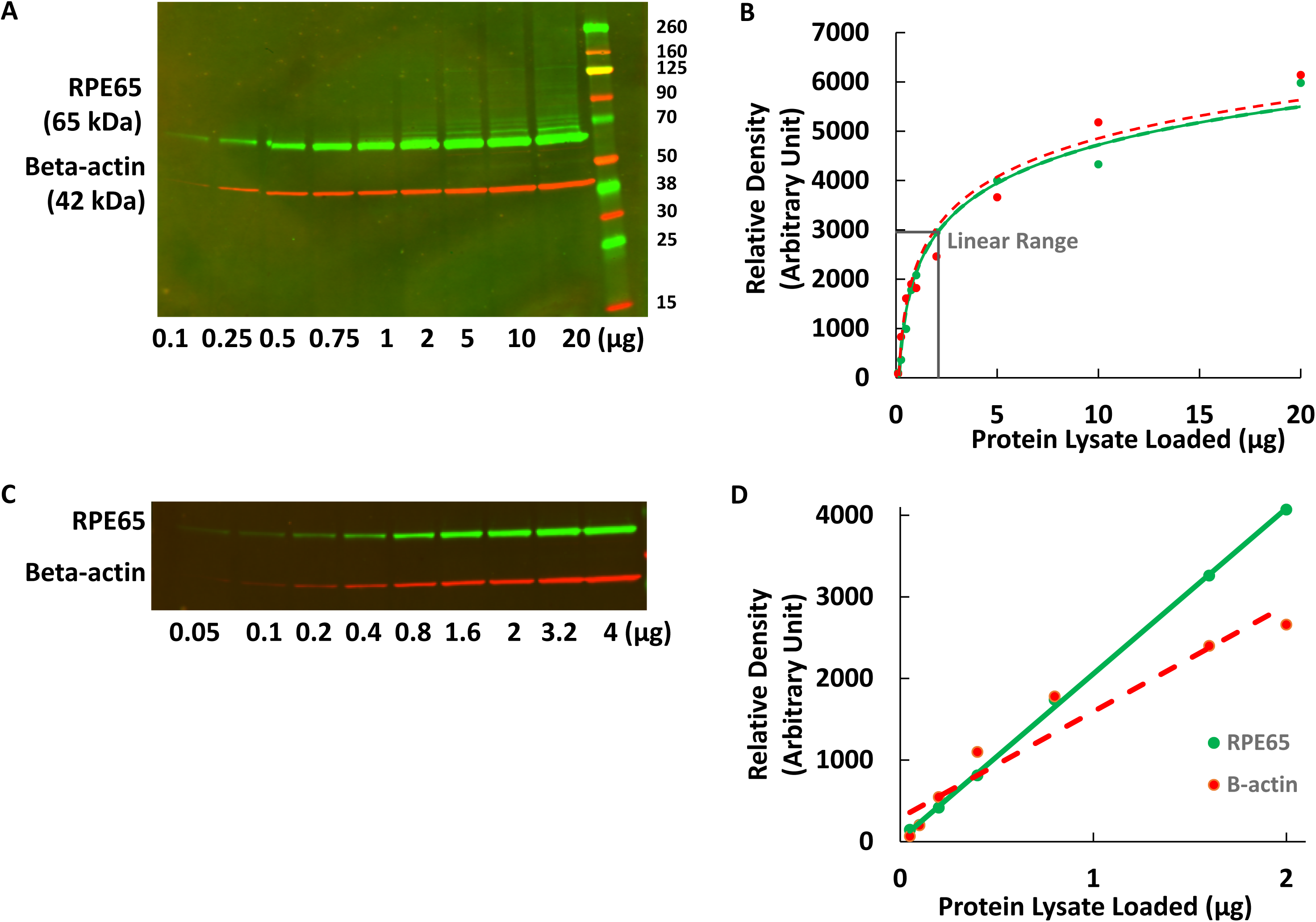
Linear range determination for RPE65 and ß-actin by Western blotting (A: 0.1 – 20 μg lysate, C: 0.05-4 μg lysate), with corresponding densitometry results (B,D). For D, the intensity was linearly correlated with protein loading amount for RPE65 (r=0.99) and for ß-actin (r=0.96).

**Figure 2.**
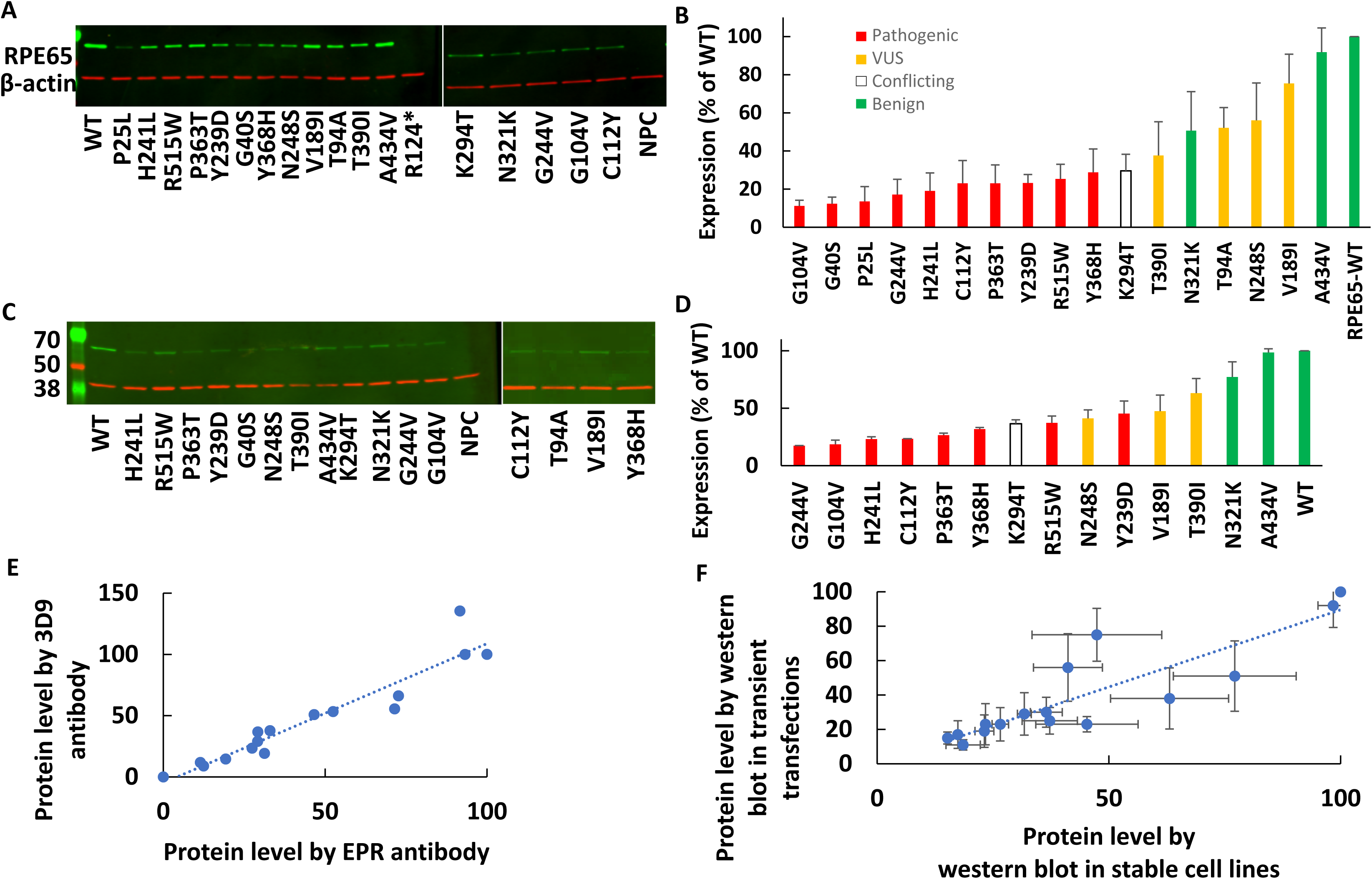
Protein expression levels of different RPE65 variants were assayed by Western blotting (A: transient transfections, C: stable cells lines) with corresponding densitometry results (B,D). In panels B and D, expression levels were normalized to ß-actin levels and are expressed as a percentage of wildtype levels (N=2-3), with known mutants shown in red, VUS in yellow, benign variants in green, and a variant with conflicting interpretations in white. E: A high correlation was observed between the expression levels measured using 2 different RPE65 antibodies (r=0.95, N=1). F: A high correlation was observed between the average measurement of RPE65 variants expressed by transient transfection and by stably-expressing cell lines (r=0.89, N=2).

#### Measurement of the protein levels of different variants by Western blot in transient transfections and stable cell lines

To analyze the effects of different variants on protein level of *RPE65* different variants (benign positive controls, pathogenic negative controls, and VUS), Western blotting was used to quantify protein levels in lysates from transient transfections (Figure 2A and B) and stable cell lines (**Figure 2C and D**). Using either transient transfection or stable cell lines, all of the pathogenic variants showed <50% of the wildtype protein level. **Figure 2E** shows a high correlation between staining of different variants with 2 different anti-*RPE65* antibodies (r=0.95), giving evidence for the specificity of the detected antigens. Also, the results showed a high correlation between measured protein levels in transiently transfected cells and in stable cell lines (r=0.88, **Figure 2F**).

#### RPE65 protein levels of different variants using flow cytometry

Flow cytometry was optimized to measure the protein levels of different *RPE65* variants using transient transfections and stable cell lines. Although the EPR antibody was best for Western blotting, for flow cytometry, antibody 3D9 showed a wider dynamic range than EPR for measuring the protein level across different variants (not shown). 3D9 was used for further staining for flow cytometry (**Figure 3**). Flow cytometry results using transiently transfected cells and stable cell lines were highly correlated (Figure 3E) (r=0.89, N=3).

**Figure 3.**
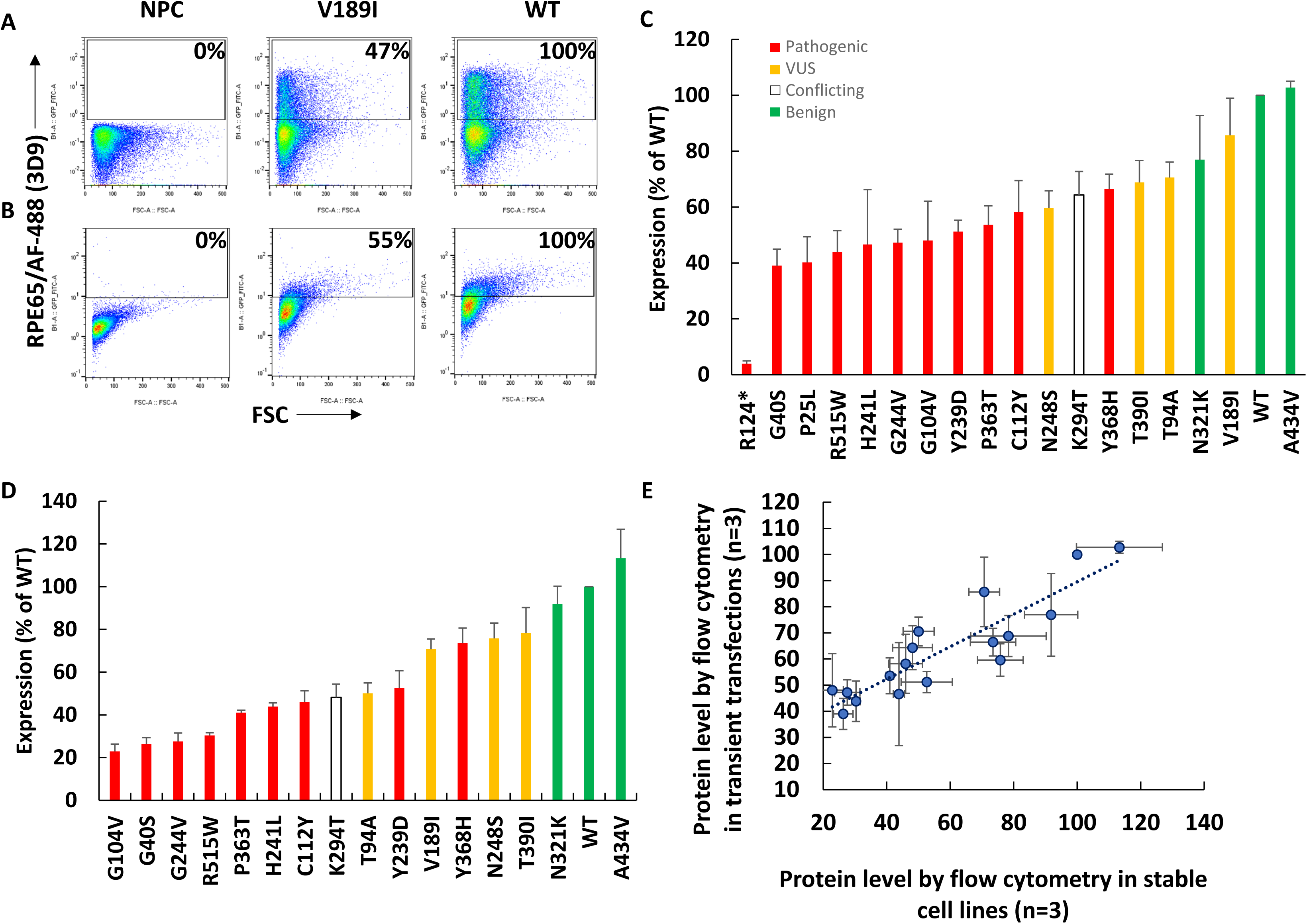
Measurement by flow cytometry of protein levels of different variants in transient transfections (A,C) and stable cell lines (B, D). Flow cytometry dot plots of untransfected cells (NPC), V189I, and wildtype RPE65 in transiently transfected cells (A) and stable cell lines (B). The transiently transfected cells express higher antigen levels than the stable cells lines with single integrations. Quantification of protein level of different variants (red: pathogenic, yellow: VUS, green: benign, white: conflicting) in transiently transfected cells (N=3). (C) and stable cell lines (D) (N=3). E: High correlation in the measurement of protein levels by flow cytometry between transiently transfected cells and stable cell lines (r=0.89, N=3).

#### Correlations between assays

Using Western blotting for quantifying protein levels in cells is not easily scalable for measuring large numbers of different variant protein expression levels. A flow cytometry assay was optimized for measuring different variants and the method showed high correlation to values measured by Western blotting using transiently transfected cells (**Figure 4A**, r=0.91) and stable cell lines (**Figure 4B**, r=0.91). Next, a pooled assay was developed for a higher throughput flow cytometry assay based on pooled RPE65 stable cell lines (**Figure 5A)**. A strong correlation was observed in the RPE65 variant expression levels between the unpooled assay and the higher-throughput pooled assay (**Figure 5B**) (r=0.87).

**Figure 4.**
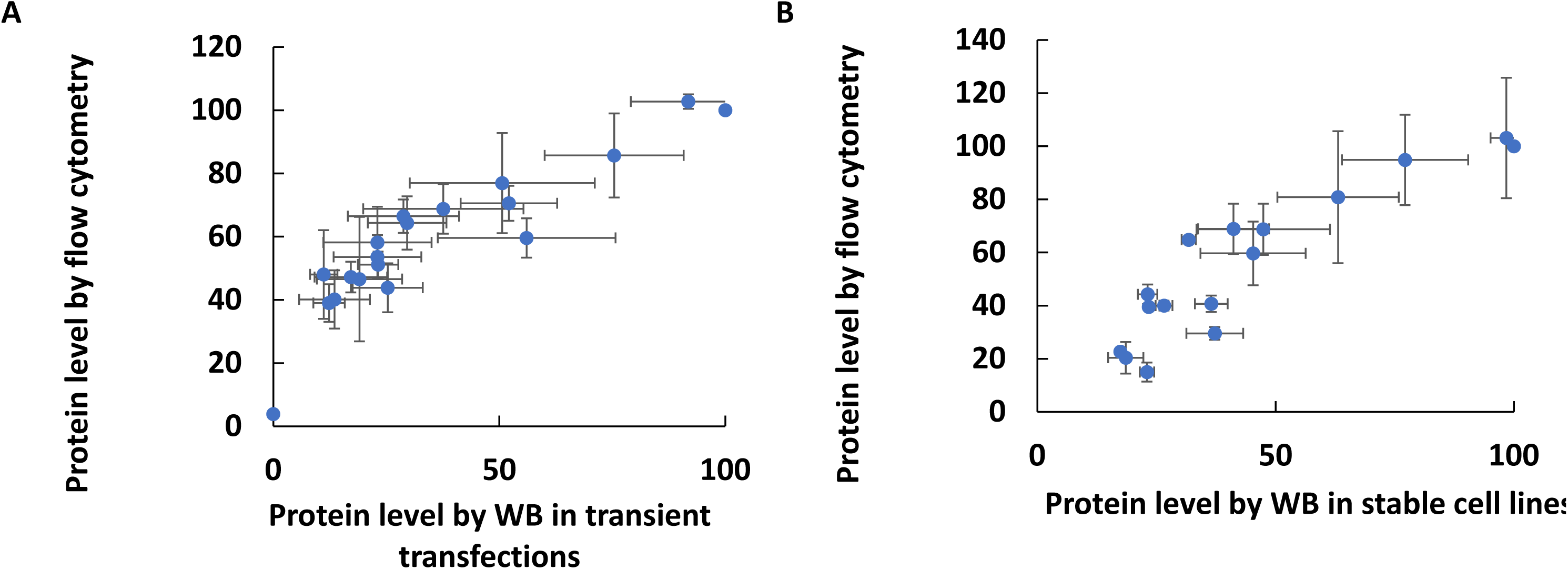
Optimized measurement of RPE65 variant protein expression levels using Western blotting with EPR antibody compared to flow cytometry with 3D9 antibody. A high correlation was obtained in both transiently transfected cells in panel A (r=0.91, N=3) and stable cell lines in panel B (r=0.91, N=2).

**Figure 5.**
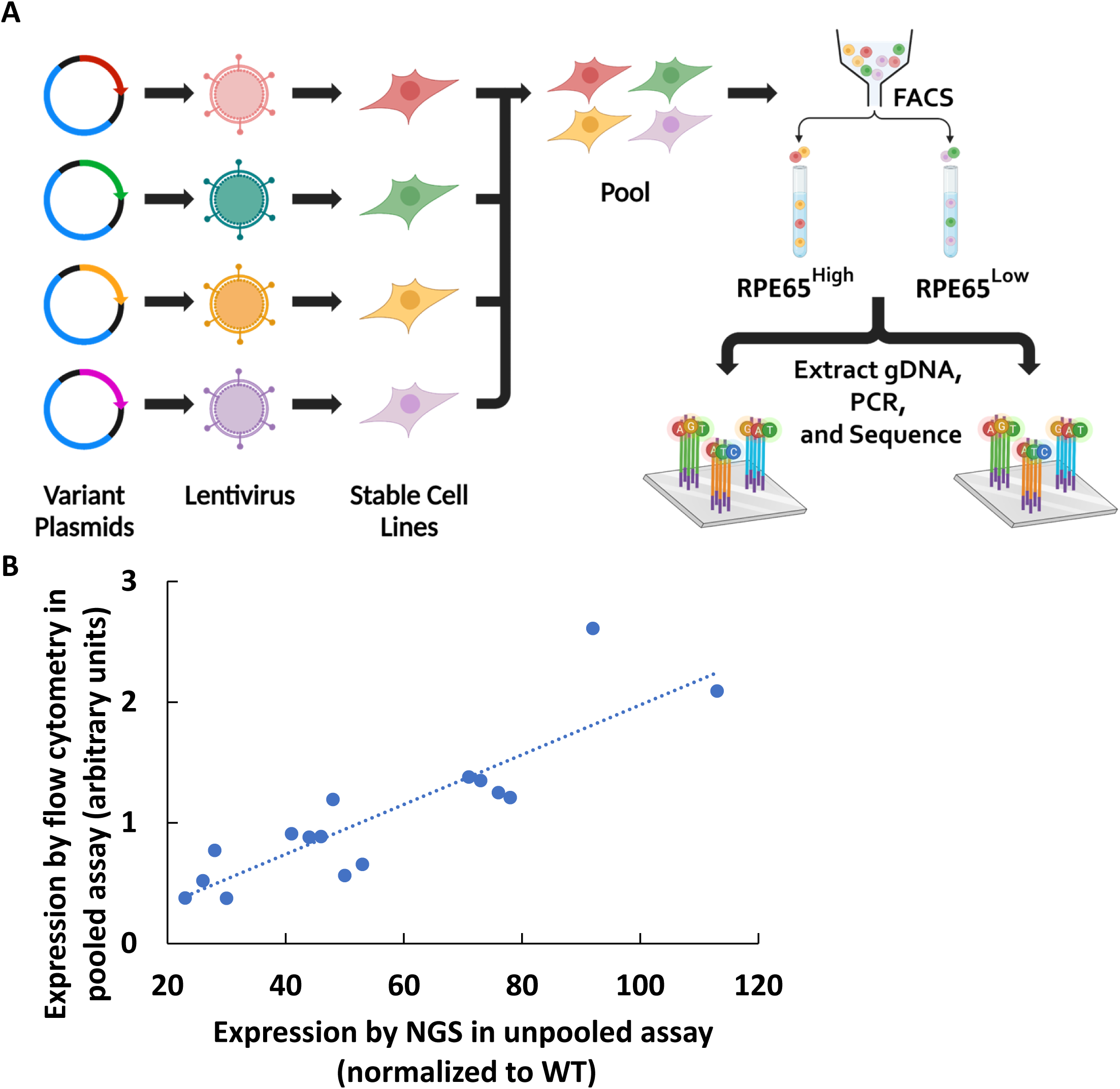
A: Workflow of pooled library assay, B: Comparing the measurement of expression levels in pooled library by NGS and flow cytometry (unpooled). There is a high correlation between pooled assay and un pooled flow cytometry assay (r=0.87, N=3)

In summary, the five assays tested above showed good agreement, as shown graphically in **Figure 6**, validating and giving support to the specificity and dynamic ranges of the assays tested above. The rare outlier points are also discussed below.

**Figure 6.**
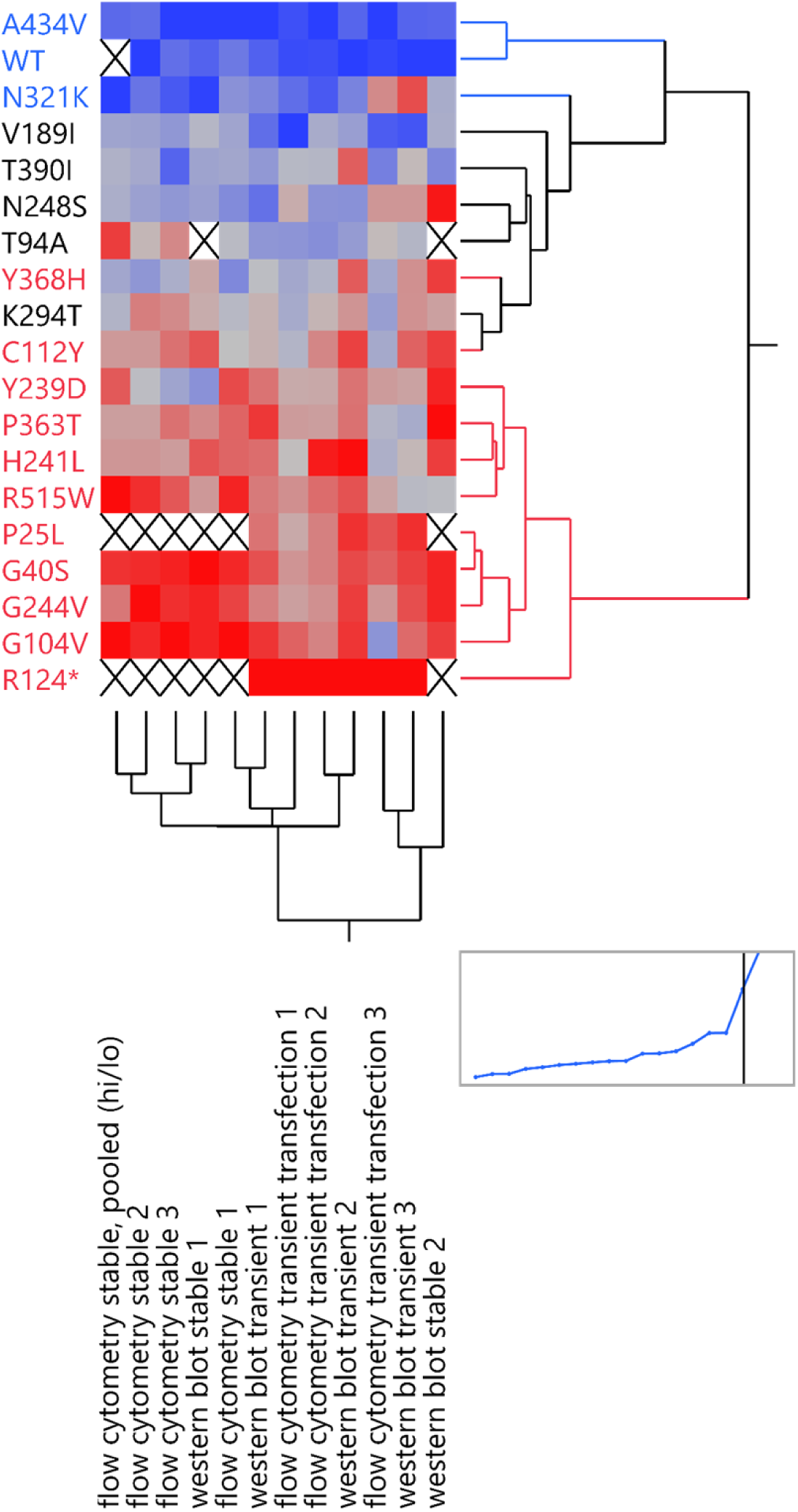
A heatmap comparing assay results across different assays (columns) and different RPE65 variants (rows). A hierarchical clustering algorithm was used to group together similar data and produce row and column dendrograms. Blue denotes high expression and red denotes low expression. An X indicates missing data.

#### Additional variants tested

Next, based on the validation of the assays shown above, additional *RPE65* variants (see Table 2 and Methods) were assayed for protein expression levels using transient transfection, staining with the 3D9 antibody, and the unpooled flow cytometry readout. **Figure 7** shows the expression levels of these additional variants. There was substantial but not complete separation between the values obtained by the positive controls (green) and the negative controls (red), as discussed below. P111S, initially a VUS, was identified as having pathogenic expression levels. Three positive control synonymous variants were hypothesized to have the same expression as the wildtype protein, but two had T385= and A434= had slightly decreased levels (71+/-5.23 % and 77+/-6.5 %, respectively), in the context of their expression from a cDNA without introns.

**Figure 7:**
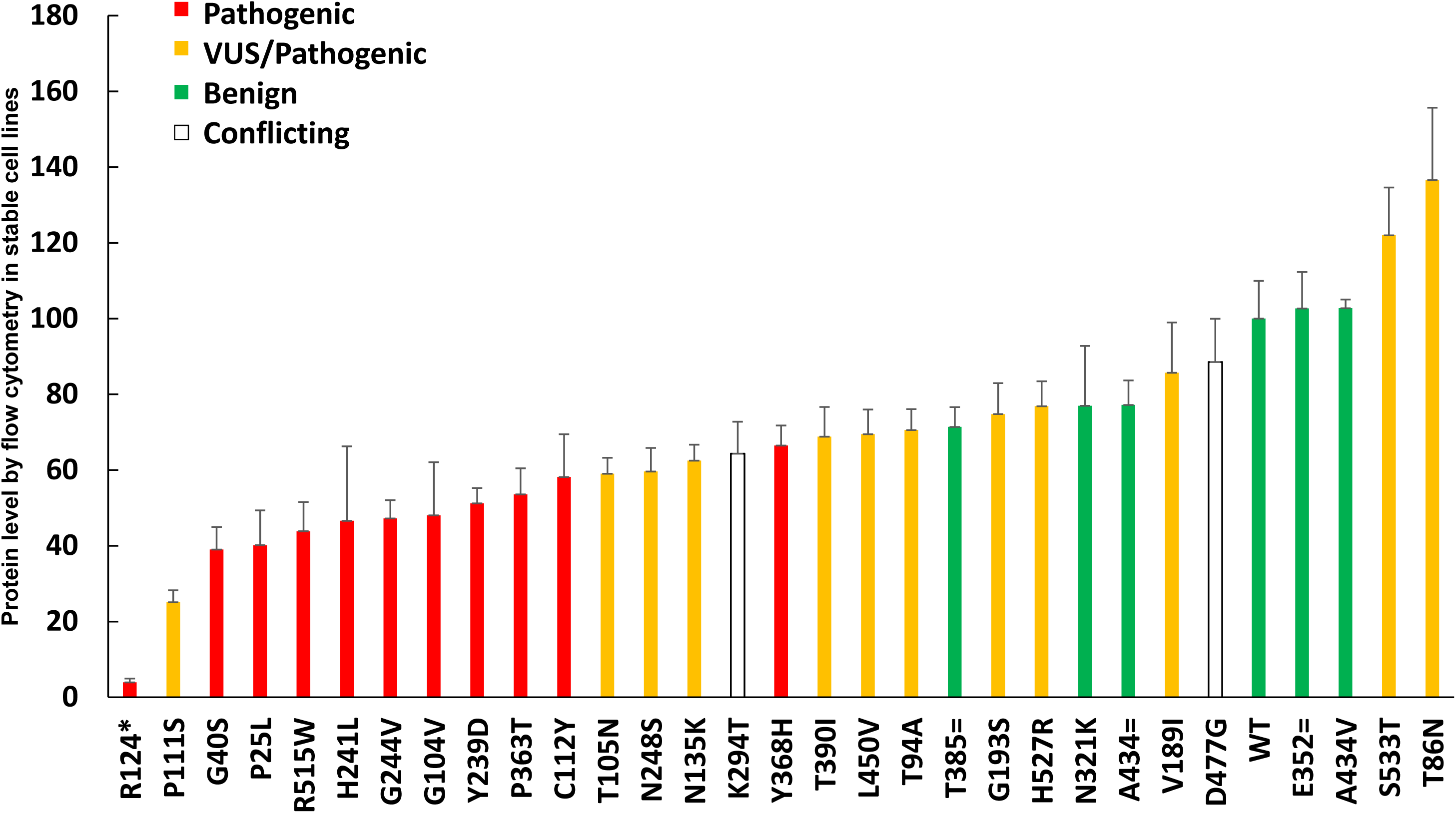
Protein levels of different RPE65 variants, measured by flow cytometry after transient transfection, N=3 (red: pathogenic, yellow: VUS, green: benign)

## Discussion

With the approval of *RPE65* gene therapy and the increased number of inherited retinal disease patients undergoing DNA sequencing, there is an increasing need for higher throughput pathogenicity assays for *RPE65* variants. The development of a higher throughput assay is often based on a simple but robust technique, and this study evaluated a straightforward protein detection assay to identify *RPE65* variants with pathogenic protein expression levels.

### Comparing the protein level measured in this study with the other studies

By one estimate, 80% of mutations are misfolding, 10% are active site mutations, and 10% have another mechanism. ^43^ Of course, these proportions and categories can vary by gene, and remain to be fully defined for *RPE65*. Based on current understanding, the different mechanisms that can be involved in pathogenicity of different *RPE65* variants include: misfolding, loss of catalytic activity, toxic gain of function, mis-localization, or aggregation of the mutated protein.

Regarding variants that are thought to be at or near the active site, we analyzed the expression level of 3 active site variants – H241L, H527R and Y239D. The mechanism of pathogenicity in the active site is different than elsewhere ^31^, but interestingly our study showed that the protein levels of H241L and Y239D are also low. In fact every pathogenic variant tested by Western blot showed an expression level of 29% of wildtype or lower. The missense H527R variant was originally reported as a VUS in ClinVar and observed in individual(s) with clinical features of *RPE65*-related conditions, but has since been reclassified as pathogenic.^44^ *RPE65* with H527R mutation did not show any catalytic activity.^31^ This study show a slightly decreased expression level (77%, Figure 7). Another variant close to active site cavity, Y239D, showed a pathogenic expression level by Western blot and flow cytometry (Table 1). However, most disease-associated missense mutations in *RPE65* are non-active site mutations.^45^

Regarding the dominant variant D477G, this variant showed wildtype expression levels in HEK293T cells, consistent with a past study showing normal expression, localization, and catalytic activity in NIH3T3 cells.^46^ More recent studies have showed a decrease of expression levels in a knock-in mouse model ^47^ and retinal degeneration only when exposed to environmental light.^48^

Regarding variants that are thought to cause misfolding and rapid degeneration of misfolded protein, including G40S, R515W, G104V and G244V, showed significantly decreased protein levels compared to wildtype (less than or equal to 29% of wildtype by Western blot). ^27,28,30,49,50^ Also, mouse models of the P25L and R91W mutations showed decreased protein levels of the mutants. ^51,52^ This decrease in protein level is likely biologically relevant; the catalytic rate of *RPE65* is so low that high expression of the protein plays the compensatory role for its weak enzymatic activity. ^21,23^ The high stability of *RPE65* with half-life >10 hours and low degradation rate lead to high abundance in the retinal pigment epithelium. ^53^

Regarding variants that may be benign, K294T was initially shown to have a severely decreased activity level. ^23^ However, as seen in dbSNP (rs61752901; accessed 12/25) a high allele fraction in Latino populations of 1.1-3.1% indicates that it is benign. The K294T variant was reported in the heterozygous state in LCA patients.^30^ Unfortunately, both a later study ^30^ and this study found borderline expression or activity levels. While we believe this variant is benign, the borderline levels do not allow that to be demonstrated definitively on a biochemical level and cannot rule out that it is hypomorphic. The other two benign variants in this study, A434V and N321K, showed normal expression.

Regarding variants that are in the amphipathic α-helix membrane targeting motif (aa 107-125), this study evaluated C112Y and P111S, which both showed pathogenic expression levels, with P111S showing the lowest expression of any missense variant tested in this study (Figure 7). Residue C112 (studied as C112A by Uppal et al) plays an important role in palmitoylation and localization of the protein into the membrane.^54,55^

### Interpreting the meaning of RPE65 expression levels

The usual mechanism causing decreased protein levels of the pathogenic *RPE65* variants is proteasomal degradation of the misfolded protein.^31^ In this study and others, the relative tendency of a particular protein sequence to misfold should be proportional in heterologous cells compared to the disease target cell, although this was not formally tested in this study. The protein detection assay in this study was successfully scaled up to a pooled assay, which in future studies can be applied efficiently to a much larger number of *RPE65* variants. However, while the absence of protein expression is good evidence of a pathogenic mutation (and can provide immediately useful information in that case), a wildtype-like expression level does not guarantee that a variant is benign. Our laboratory has been exploring the development of a higher throughput *RPE65* activity assay, intended to detect “active site” or other mutations which would affect enzyme activity but not protein levels. Despite trying to include such mutations in the panel tested, Table 1 and Figure 6 show how the protein detection assay mirrors known activity levels surprisingly well. As the number of variants tested increases, however, it would be expected to find a small fraction of “false negative” results using a protein detection assay alone.

Table 1 summarizes the protein level of different variants in previous studies compared to the current study. Both Table 1 and Figure 6 show a broad agreement between both the literature values and the assays used in this study, and among the different antibodies and techniques used. This lends higher confidence to the assays’ specificity and accuracy. The most robust and sensitive assay was Western blotting using transient transfection, and the low expression levels of the stable cell lines were limited by the single copy transgene and the presumed sensitivity of the antibody. In the pooled flow cytometry assay, the quantitative nature of counting the cells in the high versus low sorting gates, all in the same tube, is likely to increase the internal consistency of the assay compared to unpooled assays performed on separately stained and analyzed samples. While thanks to these advantages the pooled assay gave good results (Figure 6), the preferred antibody for flow cytometry, 3D9, had a moderate brightness / staining index; the dynamic range of the assay could be improved by developing a brighter antibody or, less ideally, using an epitope tag.

While the assays in this study generally had broad agreement, one specific variants (T94A) showed some disagreement between assays (Figure 6). Specifically, T94A showed borderline levels in all unpooled assays but a low level in the pooled flow cytometry assay. Of note, the T94A stable cell line did not grow well, indicating a possible toxicity of transgene expression. Selection of unusual clones during the stable cell derivation could produce artifactual results, though this does not apply to transient transfections.

Regarding the determination of an expression level which should be considered pathogenic, the pathogenic mutants showed less than or equal to 30% of protein level compared to wild type by Western blot of transiently transfected cells (Table 1) and consistent with Western blot results from other studies.^30,53^ Though the flow cytometry and Western blot results show a very high correlation (Figure 4), the Western blot values are likely more accurate on an absolute scale when comparing to external studies. By flow cytometry of transiently transfected cells, the dynamic range of the assay was slightly compressed, and less than 60% of wild type expression was always pathogenic (Table 1 and Figure 7), with 60-70% in an intermediate zone, and >70% always benign. In summary, pathogenic could be considered <30% of wild type by Western blot and <60% of wildtype by flow cytometry after transient transfection, when using the protocols of this study. While these simple numerical cutoffs are easy to understand and apply, as is often the case, many such assays have an indeterminate region or “grey area”.^56^ Ideally, the probability of pathogenicity would be expressed as a quantitative (Bayesian) probability,^57^ but for *RPE65* this will require data across a larger number of variants. Adding data from an *RPE65* activity assay would be beneficial as well.

### Reclassifying variants of unknown significance

Using the above criteria, P111S is solidly in the pathogenic range of expression levels. T105N, N248S, and N135K have borderline expression levels that may be pathogenic. V189I, S533T, and T86N show wildtype expression levels, but without an activity assay, no strong conclusions can be made.

In conclusion, all pathogenic *RPE65* variants tested show a low protein level, and validated protein expression assays can be used to reclassify the pathogenicity of VUS.^58^ Activity and localization assays would additionally identify the rare variants that have normal expression levels but lack activity.

Future work may include using the pooled assay to assay hundreds or thousands of variants. This generation of functional data will aid in the diagnosis and treatment of patients with *RPE65*-associated retinal degeneration. Furthermore, developing formal rules for the use of this functional data for classifying the pathogenicity of variants is ongoing in the Leber congenital amaurosis / early onset retinal dystrophy Variant Curation Expert Panel (LCA/eoRD VCEP) sponsored by NIH/NIGMS.

## Acknowledgements

We would like to thank T. Michael Redmond and Sheetal Uppal, Laboratory of Retinal Cell and Molecular Biology, National Eye Institute, National Institutes of Health, Bethesda, MD, USA for technical support and the gift of an RPE65 antibody. We would also like to thank Juerg Straubhaar for technical assistance, Lori Sullivan and the *RPE65* VCEP for sharing variants of interest classified as VUS, David Rousso and Spark Therapeutics for sharing a list of *RPE65* variants detected in patients, and the Genomics Core of the Ocular Genomics Institute for next-generation sequencing. Xiaoping Yang (Broad Institute, Cambridge, MA) designed the pMT025 primer sequences used in this study.

---

**Supplementary table 1.**
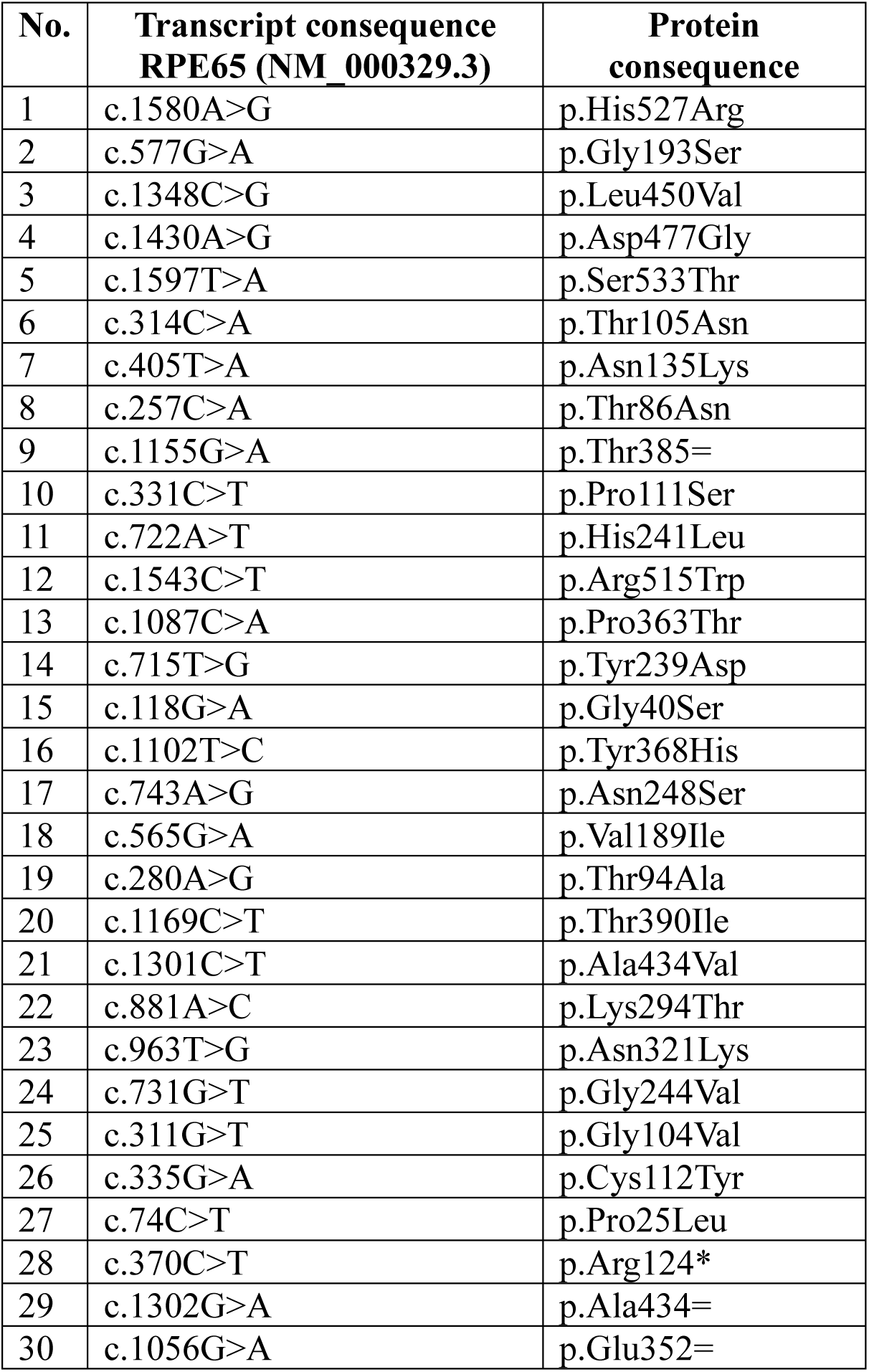
List of different RPE65 variants tested in this study with HGVS nomenclature. The reference transcript is NM_000329.3.

**Supplementary Table 2.** Literature review of expression levels and catalytic activities of additional RPE65 variants.

| No. | Variants | Pathogenicity | Literature expression % | Literature activity % | References |
| --- | --- | --- | --- | --- | --- |
| 1 | Thr101Ile | Pathogenic | ~100, ~15 | 1.27, 2.4 | 30,31 |
| 2 | Leu408Pro | Pathogenic | ~80, ~20 | 5.86, 5 | 30,31 |
| 3 | Tyr79His | Pathogenic | 3.31 | 2.5 | 25 |
| 4 | Glu95Gln | Pathogenic (mild) | 28.07 | 6.1 | 25 |
| 5 | Leu22Pro | Pathogenic | 32.4, ~20 | 13.5, 3.9 | 25,31 |
| 6 | Arg44Gln | Pathogenic | ~75, ~25 | <2, 1.34 | 23,30,31 |
| 7 | Arg91Gln | Pathogenic | ~110 | 1.33 | 30 |
| 8 | His68Tyr | Pathogenic |  | <2 | 23 |
| 9 | Ala132Thr | Pathogenic |  | 50 | 23 |
| 10 | His182Tyr | Pathogenic |  | 10 | 23 |
| 11 | Glu417Gln | Pathogenic |  | <2 | 23 |
| 12 | Gly528Val | Pathogenic |  | <2 | 23 |
| 13 | Cys330Tyr | Pathogenic | 0 | <2, 1.6, 1.6 | 23,31,45 |
| 14 | Ala434Val | Benign |  | 55 | 23 |
| 15 | Glu417Gln | Pathogenic | ~75 | 1.1 | 26,31 |
| 16 | His313Arg | Pathogenic |  | Not Detectable | 31 |
| 17 | Tyr318Asn | Pathogenic | ~20, ~20 | 5.12, 7.1 | 30,31 |
| 18 | Arg91Trp | Pathogenic | ~10 | 5.08 | 30 |

## Supplemental Methods

### Generation of stable cell line for *RPE65* variants

For lentiviral production, HEK293FT (Thermo Fisher, Cat no. R70007) cells were seeded at a density of 0.4×10^6^ cells/well of 6-well plate. Cloned pMT_025 *RPE65* lentiviral expression plasmid was co-transfected with pMD2.G (Addgene no. 12259) and psPAX2 (Addgene no. 12260) using Lipofectamine LTX transfection reagent (Thermo Fisher, Cat no. 1533803) following the manufacturer’s instructions. Fresh media was supplemented after 24 hours and viral supernatant was harvested at 48, 72 and 96 hours, pooled, filtered and concentrated by using Lenti-X-Concentrator (Takara, Cat no.631232) according to the manufacturer’s instructions. Concentrated aliquots were frozen in − 80 °C until use. Titers were determined by transduction in HEK293T cells followed by Crystal Violet staining. For the generation of stable lines, 1×10^5^ HEK-293T cells were plated in a well of 6-well plate and transduced at an MOI of 0.3 by spinfection (2000g for 20 minutes) in the presence of Polybrene (8 µg/mL, Sigma, Cat no.107689). Media was replaced the following day. On day 3, puromycin (600 ng/mL) was added, selected for five days, and then expanded to establish stable lines. To confirm the integration, cells were lysed using Quick Extract Buffer (Lucigen, Cat no. QE09050), and PCR was performed using Q5 Master Mix (New England Biolabs, Cat no. M0492S) using primers XY304 (ATTCTCCTTGGAATTTGCCCTTT) and XY305 (CATAGCGTAAAAGGAGCAACA).

### Quantification of *RPE65* levels using Western blots

**Cell** lysates were prepared for quantitating RPE65 protein levels 48 hours post-transfections. For stable cells lines, lysates were prepared at 80% confluency. Briefly, the cells were washed with PBS, trypsinized and washed again with cold PBS, and lysed using cold 1X RIPA buffer (Abcam, Cat no. ab156034) supplemented with protease inhibitor cocktails (Roche, Cat no. 11697498001). Lysates were agitated at 4°C for 30 minutes and by briefly vortexing for 15 seconds, periodically every 10 minutes. Lysates were centrifuged at 18000×g for 30 minutes at 4°C and the supernatant was collected and stored at −80°C until use. Few variants (X, Y, Z) showed reduced growth rates in culture and, therefore, were excluded from the analysis (N=?)

Protein concentration was determined using Pierce BCA protein assay kit (Thermo Fisher Scientific, Cat no. 23225) according to the manufacturer’s protocol. To assure correct quantitative immunoblots of *RPE65* different variants, different concentrations of whole cell lysate (0.1-20 μg) from wildtype *RPE65*-expressing cells were loaded onto the SDS-PAGE (4-20% tris-glycine gel, Invitrogen Cat no XP04200BOX) to find the linear range where the total protein loaded was linearly related to fluorescent band intensity. For electrophoresis loading, each sample was mixed with 4X LDS sample loading buffer (Invitrogen, Cat no. NP0007), denatured by heating at 70°C for 10 min, spun for 1 min at 18000×g and the supernatants were loaded onto the SDS-PAGE gel. Gel electrophoresis was performed at 150V for ∼90 minutes.

Then proteins were transferred from the gels onto polyvinylidene difluoride (PVDF) blotting membranes using the iBlot™ 2 system (Invitrogen, Carlsbad, CA) for 20V for 1 minute, 23V for 4 minutes and 25V for 2 minutes. Total protein staining was performed using Revert 700 following manufacturer’s protocol (LICOR, Cat no.926-11011). For staining with *RPE65* and ß-actin antibodies, the membranes were blocked for 1 hour using Intercept® blocking buffer (LICOR, Cat no.927-60001), then incubated with 1:2500 or 1:5000 dilution of *RPE65* primary antibody and 1:5000 dilution of ß-actin primary antibody overnight at 4°C. The primary antibodies for *RPE65* were rabbit monoclonal [EPR7024(N)] anti-*RPE65* C-terminal (Abcam, ab175936), herein referred to as “EPR”, and mouse monoclonal anti-*RPE65* (401.8B11.3D9), (Novus Biologicals, Cat no. NB100-355), herein referred to as “3D9”. For ß-actin, mouse monoclonal IgG1 (Santa Cruz biotechnology, Cat no. sc-47778) and rabbit polyclonal (Abcam, Cat no. ab8227) were used. The following day, the membranes were washed 3 times with TBST (Tris buffered saline with 0.1% Tween-20) for 5 minutes and incubated with a 1:10,000 dilution of IRDye® 800CW and 700CW secondary antibodies for 1 hour followed by three 5 min washes in TBST. The secondary antibodies were IRDye 680RD Goat anti-Mouse (LICOR, Cat no. 926-68070) and IRDye 800CW Goat anti-Rabbit (Licor, Cat no. 926-32211). The membranes were then imaged on the 800-nm and 700-nm wavelength channel using the Odyssey CLx infrared imaging system (LICOR, Lincoln, Nebraska). The density of Western blot bands was measured using Odyssey software and normalized to ß-actin internal protein levels. After determining the proportional linear range for detection of ß-actin and *RPE65*, Western blots of lysates from *RPE65* transient transfection and stable cell lines were performed using 1µg of total protein for transient transfections and 4µg of total protein for stable cell lines, as described above. All data were then normalized to the expression level seen in the wild-type (WT) *RPE65* sample.

